# Development of Audiovisual Integration in Beginning Readers: A Longitudinal fMRI Study

**DOI:** 10.64898/2026.07.19.739407

**Authors:** Dalia Braverman-Jaiven, Rola Farah, Dror Kraus, Ori Zehngut, Tomer Michaeli, Roi Carmel, Ayelet Elor, Mika Shapira Rootman, Michael A. Skeide, Joanna Finnemann, Tzipi Horowitz-Kraus

## Abstract

Acquiring reading proficiency, unlike spoken language, requires the brain to engage several specialized neural systems to map visual symbols (letters) to their corresponding phonological sounds (audiovisual integration), laying the foundation for fluent reading.

This study investigates the neural and behavioral trajectory of developing audiovisual (AV) integration during the first year of learning to read. Thirty-two healthy Hebrew-speaking first-grade children were assessed at 3 time points across the school year: beginning, middle, and end of first grade. Participants underwent behavioral testing and brain fMRI scans while performing a block-design fMRI task involving AV matching or non-matching letters and sounds.

Together with improvement in behavioral test scores related to general abilities, working memory, cognitive flexibility, and phonemic abilities, a Drift Diffusion Modeling (DDM) analysis of the accuracy and reaction time across sessions suggested faster and more efficient responses as the year progressed. fMRI results showed a significant increase in activation, from the beginning to the end of the first grade, in the left superior temporal gyrus (STG), frontal cortices and parietotemporal cortices, with a shift towards left-lateralization in the fusiform gyrus at the end of the year. These findings point towards the second half of the first grade as the time window for neural specialization and lateralization to phonological and orthographic information. A significant positive correlation between the fusiform gyrus activation and naming objects and colors scores across all sessions links this neural specialization to cognitive flexibility behavioral skills.

**Key points:**

- Significant increase, predominantly between the middle and end of the first grade, in activation in fusiform, left superior temporal, frontal, and parietotemporal cortices across the first grade.
- Significant left lateralization in the fusiform gyrus at the end of first grade.
- Positive correlation between naming skills and a bilateral fusiform gyrus activation throughout the first grade.

## Introduction

### The neurobiology of reading

Learning to read is an effortful process that requires the brain to engage in sensory and cognitive information processing. This process is different from learning to speak or talk, probably due to reading being a recent ability that human brains are not naturally shaped to learn (Holloway et al., 2015). Frith describes that the process of learning to read involves three basic strategies that a child has to master to become a fluent reader: logographic, alphabetic and orthographic (Frith, 1986). The logographic refers to the initial recognition of a word based on graphic features only, meaning the ability of the child to recognize a logo, for example, based on its shape but without being able to actually read the word. The alphabet strategy refers to recognizing that each letter represents a sound and being able to put these sounds together to evoke a word. The third and final strategy, orthographic, refers to instantly recognizing a word, taking into account the letter order but not the letter sound (Frith, 1986).

Since reading is initially very cognitively demanding, it does not rely on a single brain region or network, but rather recruits several neural systems, primarily in the left hemisphere, along with executive functions (EF) and attention. fMRI studies reveal three primary neural systems involved in reading: The anterior system (inferior frontal gyrus and Broca’s area) (Shaywitz & Shaywitz, 2007) associated with phonological processing and syntactic processing and responsible for articulation, word analysis and the production of language (Joseph et al., 2001). The parieto-temporal system (angular gyrus and supramarginal gyrus) is associated with mapping visual input into sound (Kweldju, 2015) and is critical for the decomposition of words into phonemes (Joseph et al., 2001). The occipito-temporal system acts as the visual word form area (VWFA) (Kweldju, 2015; Shaywitz & Shaywitz, 2007), specialized in rapid and automatic identification of words (Joseph et al., 2001).

Other processes and cognitive abilities, such as audiovisual (AV) integration, attention, and executive functions, are crucial for reading development, therefore indirectly involving other regions of the brain. Initial visual analysis activates the extrastriate visual cortex, specifically the left medial extrastriate cortex and fusiform gyrus (Joseph et al., 2001). Phonological processing, converting letters into sounds, activates the superior temporal gyrus and the inferior frontal cortex (Joseph et al., 2001; Kweldju, 2015).

During the early stages of reading (ages 5 to 7), children typically exhibit ambilaterality, meaning both sides of the brain are utilized to process the information required (Sadick & Ginsburg, 1978). However, as reading develops and becomes automatic and fluent (children aged 8 to 11 years), there is a strong shift of activations towards the left hemisphere (Sadick & Ginsburg, 1978). This process does not happen as a result of an increase in activation in the left hemisphere, but rather a reduced activation of the right hemisphere as expertise develops (Seghier & Price, 2011).

### Audiovisual integration during reading acquisition

Audiovisual (AV) integration refers to the ability to integrate auditory and visual information accurately and effectively (Horowitz-Kraus et al., 2025). The foundation of fluent reading lies in the ability to map visual shapes (graphemes or letters) to their corresponding sounds (phonemes). This ability transitions to highly automatic word recognition and this transition relies on the brain’s ability to synchronize its visual and auditory networks rapidly (Ren et al., 2026). In the brain, the superior temporal gyrus (STG) and Heschl’s gyrus act as auditory association cortices responsible for phonological processing (Ren et al., 2026). The occipitotemporal cortex, specifically the fusiform gyrus, which houses the Visual Word Form Area (VWFA) (McCandliss et al., 2003), plays an important role in processing written characters, driven by phonological processing (Pleisch et al., 2019). This neural response develops rapidly during the first year of reading acquisition, with the activation in this region along with the STG and IFG increasing significantly in healthy developing children (Karipidis et al., 2021).

As reading develops, matching letters to sounds engages the left inferior frontal gyrus (IFG) and its connections to the STG and the ventral occipital cortex (vOTC) from pre-reading stages to reading development (Karipidis et al., 2021; Ren et al., 2026). This development is observed in typically developing children, whereas children with poor reading outcomes fail to show it (Karipidis et al., 2021). These findings converge on four functionally distinct contributions to letter-sound mapping, which together define the regions of interest in the present study. The fusiform gyrus is responsible for orthographic processing and automatic recognition of written words (Horowitz Kraus, 2023; McCandliss et al., 2003), whereas the superior temporal gyrus supports phonological processing, recognition of auditory words and auditory feedback processing (Joseph et al., 2001). These two regions provide the visual and auditory input for letter-sound matching. The frontal and parietotemporal cortices serve as the broader anatomical regions housing the IFG and the angular and supramarginal gyri, respectively, and perform this mapping precisely. The parietotemporal cortex converts graphemes into phonemes and evaluates lexical and sublexical phonology (Joseph et al., 2001; Shaywitz & Shaywitz, 2007). Finally, the frontal cortex supports articulation, word analysis, and syntactic processing (Kweldju, 2015).

While AV integration is fundamental to reading acquisition, the developmental trajectory of letter-sound congruency processing during the first year of schooling remains unclear. This study aims to track how the brain’s sensitivity to letter-sound congruency develops across the first grade in Hebrew-speaking children, as a window into the more general development of AV integration during reading acquisition.

We hypothesize that audiovisual matching performance will improve over the school year, accompanied by an increase in activation in the fusiform gyrus, superior temporal gyrus, frontal and parietotemporal cortices, involved in AV integration and letter-sound congruency, and that this increase will become left lateralized as reading skills develop. We further predict that this increase in activation will be positively correlated with behavioral naming scores, linking neural sensitivity to letter-sound congruency with behavioral processing speed and executive functions directly related to reading abilities.

## Methods

### Participants

Thirty-two Hebrew-speaking children entering the first grade were recruited for this study (age at first testing: 77.6 ± 4.55 months; 22 males and 27 right-handed) via posted ads between December 2023 and June 2025. All children had normal or corrected-to-normal vision and hearing, and parents did not report any learning difficulties or neurological disorders. Children performed several behavioral tests for general intelligence, executive functions, reading readiness, and attention, as well as a functional MRI session while performing an audiovisual matching task. All participants provided their verbal consent, and their parents signed a consent form. The study was approved by the committee and adhered to the guidelines specified in the Declaration of Helsinki.

### Study design

In this longitudinal study, participants underwent functional and structural MRI scanning and behavioral assessment at three time points: beginning, middle, and end of the first grade. Thirty-two children (age: 77.6 ± 4.55 months) completed the first assessment (Test 1; T1), twenty-eight children (age: 83.6 ± 3.77 months) completed the second assessment (Test 2; T2), and twenty-six children (age: 86.9 ± 3.90 months) completed the third assessment at the end of the school year (Test 3; T3).

### Behavioral measures

Participants underwent a battery of behavioral tests during each assessment. These tests help create a cognitive profile for each participant and quantitatively evaluate key cognitive abilities related to reading acquisition.

General intelligence and verbal abilities were measured through the Picture Naming subtest from the Wechsler Preschool and Primary Scale of Intelligence Battery (Wechsler, 2012).

Executive functions were measured in several domains using standardized tests. Processing speed was measured through the Symbol Detection subtest from the WPPSI battery (Wechsler, 2012). Working memory was evaluated through the Memory of Digits subtest from the Comprehensive Test of Phonological Processing (CTOPP) (Wagner et al., 1999). Inhibition was measured through the Walk / Don’t Walk subtest from the Test of Everyday Attention for Children (TEA-Ch) battery (Manly et al., 2001). Cognitive flexibility was measured using the Animals and colors inhibition/switching Test (Ziv, 2017) and the Naming Mixed Objects subtest from the Aleph-Taph battery (Shany et al., 2006). Auditory attention was measured through the Score! Subtest from the TEA-Ch battery (Manly et al., 2001). Selective attention was measured using the Sky-Search subtest from the TEA-Ch battery (Manly et al., 2001). Finally, reading abilities were measured through the Naming Letters subtest from the Shatil battery (Shatil, 2000; Shatil & Share, 2003).

### Neuroimaging measures

#### The Audiovisual integration fMRI task

During the fMRI scan, the children performed an audiovisual matching task involving letter-sound correspondence. The paradigm employed a block design comprising 24 blocks across 4 conditions presented in a pseudorandomized order: (1) Audiovisual (AV) match, where the visually presented letter matches the sound; (2) audio only where the sound is presented with no visual stimuli; (3) visual only, where a letter is displayed with no auditory stimuli; and (4) audiovisual (AV) non-match where the letter and sound are incongruent. Stimuli consisted of six Hebrew consonants: בּ (/b/ - “bet”), פּ (/p/ - “pe”), פ (/f/ - “fe”), כּ (/k/ - “kaf”), מ (/m/ - “mem”) and ט (/t/ - “tet”).

Each block lasted 4 seconds and consisted of 6 stimuli presented for 467 ms followed by a 200 ms inter-stimuli interval. Blocks were separated by a four-second resting period. The task structure is visualized in Figure 1. Although we employed a passive viewing and listening task requiring no overt response, children were asked to press a button upon the appearance of a target oddball stimulus (a spaceship or bell sound) to ensure sustained attention. Each condition had approximately 24 seconds of data. The tasks lasted a total of 5 minutes. The task was presented using the E-Prime 3.0 stimulus presentation program and Chronos (Psychology Software Tools, Pittsburgh, PA, USA) response collection (PSTNET, 2020). Auditory stimuli were presented using the MR-compatible OptoACTIVE noise-canceling headphones. (Optoacoustics, Israel).

**Figure 1:**
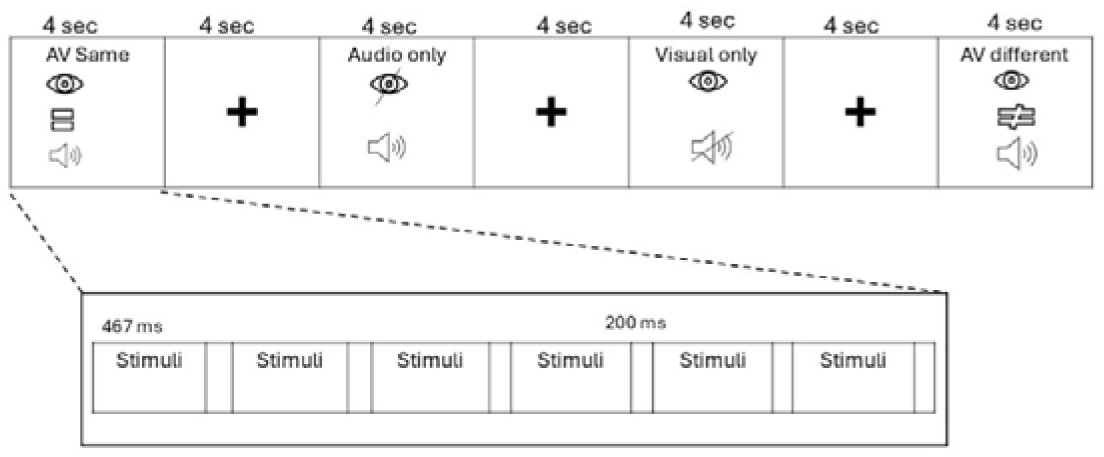
Schematic of the block-design audiovisual matching task. Each 4-second task block consists of 6 stimulus events (467 ms of stimulus presentation followed by a 200 ms inter-stimulus interval) and is followed by a 4-second rest block. The task consisted of 24 blocks including six blocks for each of the four conditions presented in a pseudorandomized order: AV same, audio, visual and AV different. AV = audiovisual.

### Data acquisition

MRI data were acquired in the May-Blum-Dahl Human MRI Research Center (TechMRC) at the Technion – Israel Institute of Technology, Israel. Prior to entering the scanner, participants practiced the task in a mock scanner to understand the procedures and to get comfortable with the environment. During this practice session, children completed the task and were to press one button for AV match trials and a second button for AV non-match trials. Accuracy and reaction times recorded in this session were later extracted and analyzed with drift diffusion modeling (DDM).

During the scan, three to four different runs of the task were performed, depending on the participant’s comfort and compliance. Scans were conducted in a Siemens Healthineers MAGNETOM Prisma 3T MRI (Siemens Healthineers, Erlangen, Germany) with a 64-channel head/neck receive coil. Anatomical images were acquired using a T1-weighted MPRAGE pulse sequence with the following parameters: Repetition time (TR): 2000 ms, Echo time (TE): 3.24 ms, Inversion time: 900 ms, Flip Angle (FA): 9 degrees, Matrix size: 256 x 240 x 176 and Voxel size: 1 mm. Functional BOLD images were collected by T2*-weighted echo planar imaging (EPI) with the following parameters: TE/TR = 22/2000 ms, Flip Angle (FA): 80 degrees, Matrix size: 82 x 82, Voxel size: 2.49 x 2.49 x 2.5 mm, Number of slices: 60, Slice thickness: 2.5 mm, Slice gap: 0.25 mm.

### Data analysis Behavioral measures

Statistical analysis was conducted using IBM SPSS (IBM) Jamovi 2.6.44 (Jamovi, 2024; Team, 2024). Descriptive statistics (mean and standard deviation) were computed to characterize the sample’s cognitive profile. Repeated-measures ANOVAs (Singmann et al., 2012) were conducted for all behavioral measures to identify differences in scores across all three sessions. Post-hoc t-tests (Lenth, 2023) were also conducted to identify effects driving significant differences.

Accuracy and reaction times were extracted when children had to respond to AV match and AV non-match stimuli. These data were analyzed using drift diffusion modeling (Myers et al., 2022) to calculate the drift rate, which indicates the speed and quality of information processing.

### MRI data processing

Anatomical and functional MRI data preprocessing was conducted using fMRIprep version 25.2.4, (Esteban et al., 2019). The preprocessing pipeline included realignment, coregistration of the functional data to the anatomical data, and normalization to the Montreal Neurological Institute (MNI) standard stereotactic space. Head motion was quantified using framewise displacement (FD) as part of the fMRIprep confound output (Power et al., 2012). The six rigid-body motion parameters, their temporal derivatives, and squared terms (24-parameter model) were included as nuisance regressors in the first-level GLM for each run. To verify that motion did not affect the longitudinal pattern of the results, mean FD was compared across sessions using a repeated-measures ANOVA and no significant difference was found (F (2,48) = 0.038, p=0.962).

The first-level statistical model was estimated using the General Linear Model (GLM) implemented in the Nilearn Python library (Abraham et al., 2014). A design matrix was constructed for each subject, modeling the onset of each stimulus condition convolved with a standard hemodynamic response function (HRF). Prior to model fitting, functional data were spatially smoothed using a 6.0 mm FWHM Gaussian kernel. Contrast maps were computed for each subject to compare responses to different stimuli, yielding normalized Z-score maps that represent the magnitude and statistical significance of activation for each voxel.

Second-level analysis was conducted to determine the group-level effects. A smoothing kernel of 8.0mm was used prior to model fitting. Because sequential Gaussian smoothing combines in quadrature, this is equivalent to an effective smoothing kernel of 10 mm FWHM at a group level. This kernel was chosen to address the lower signal-to-noise ratio typical in pediatric samples. The individual z-maps were averaged, and a one-sample t-test was performed for each contrast to assess whether the mean activation at each voxel was significantly different across all subjects.

For the Region of Interest (ROI) analysis, four anatomical masks were placed in key regions related to AV integration, including frontal cortex, parietotemporal cortex, fusiform gyrus, and superior temporal gyrus, using the Harvard-Oxford atlas through Nilearn in Python. All masks were divided into left and right hemispheres, resulting in three masks for each region (bilateral, left, and right). Given the a priori selection of ROIs based on established literature, no correction for multiple comparisons was applied across regions.

Mean beta values were extracted for the AV match > AV nonmatch contrast for each mask and session. To assess longitudinal changes, repeated-measures ANOVAs were performed to identify differences between sessions for each mask. Post-hoc paired t-tests were then conducted using the values of all three masks (bilateral, left and right). To examine the lateralization effect, mean beta values extracted from the left and right hemispheres were compared, and one-tailed paired t-tests were conducted for each time point, testing the hypothesis that left hemisphere activation is significantly greater than right hemisphere activation.

### Correlations between behavioral and neuroimaging data

The relationship between naming objects and colors standard scores and ROI activation was tested using a linear mixed effects model with naming scores and session as fixed effects and a by-subject random intercept to account for repeated measurements across the three sessions (Mean Beta ∼ Naming score + Session + (1 | Subject)).

## Results

### Behavioral results

Repeated measures ANOVA on the behavioral scores revealed that a significant increase with age was observed in verbal abilities (WPPSI Naming), working memory (digit span, CTOPP), and cognitive flexibility (Switching animals and colors and Naming Mixed Objects). In phonemic awareness (Shatil phoneme awareness task) and inhibition (Walk don’t Walk – TEA-Ch), there was a significant increase in the scores between the beginning and the middle of the school year, but the effect did not persist until the end of the year. Repeated measures ANOVAs were conducted with listwise deletion per measure, such that only participants with complete data across all three sessions for a given measure were included, resulting in varying sample sizes across analyses (Table 1).

**Table 1.**
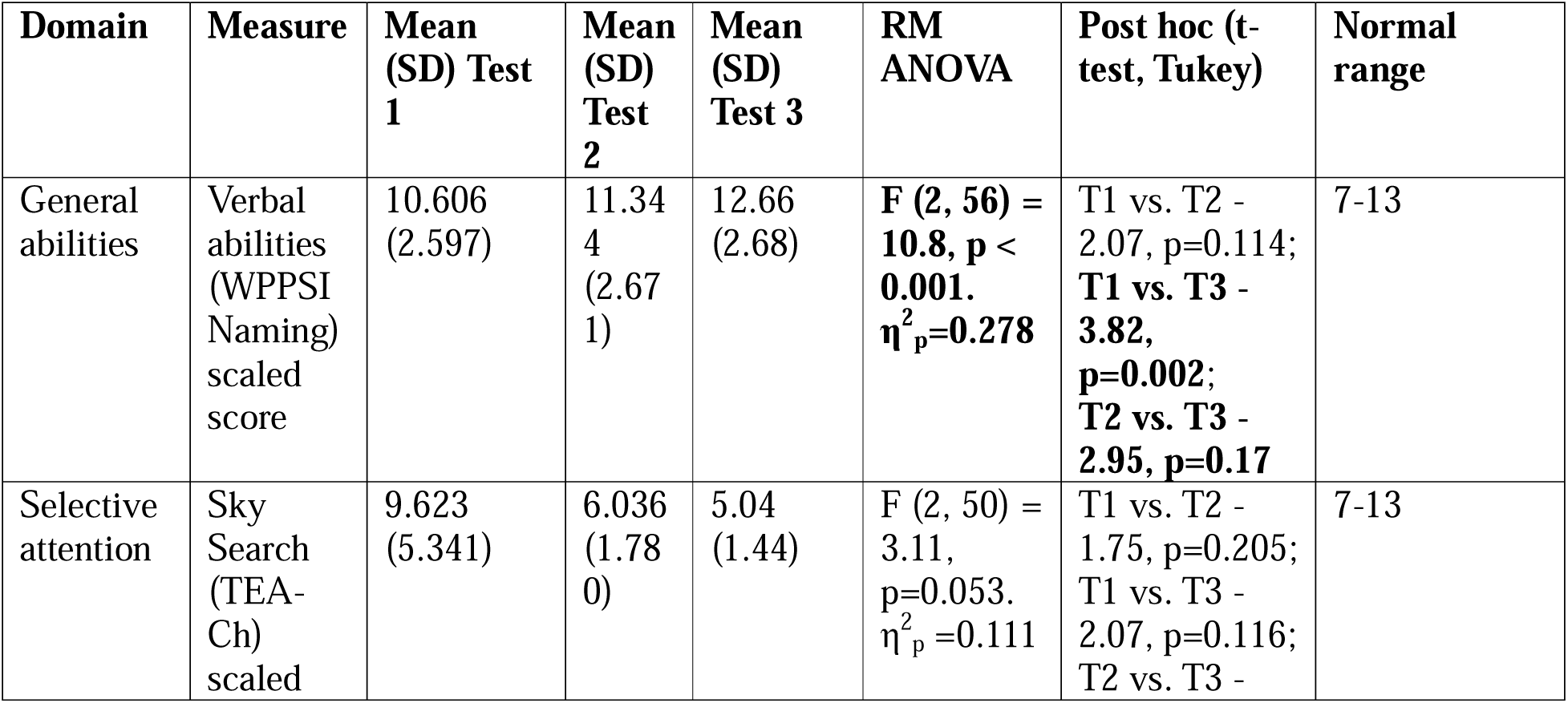
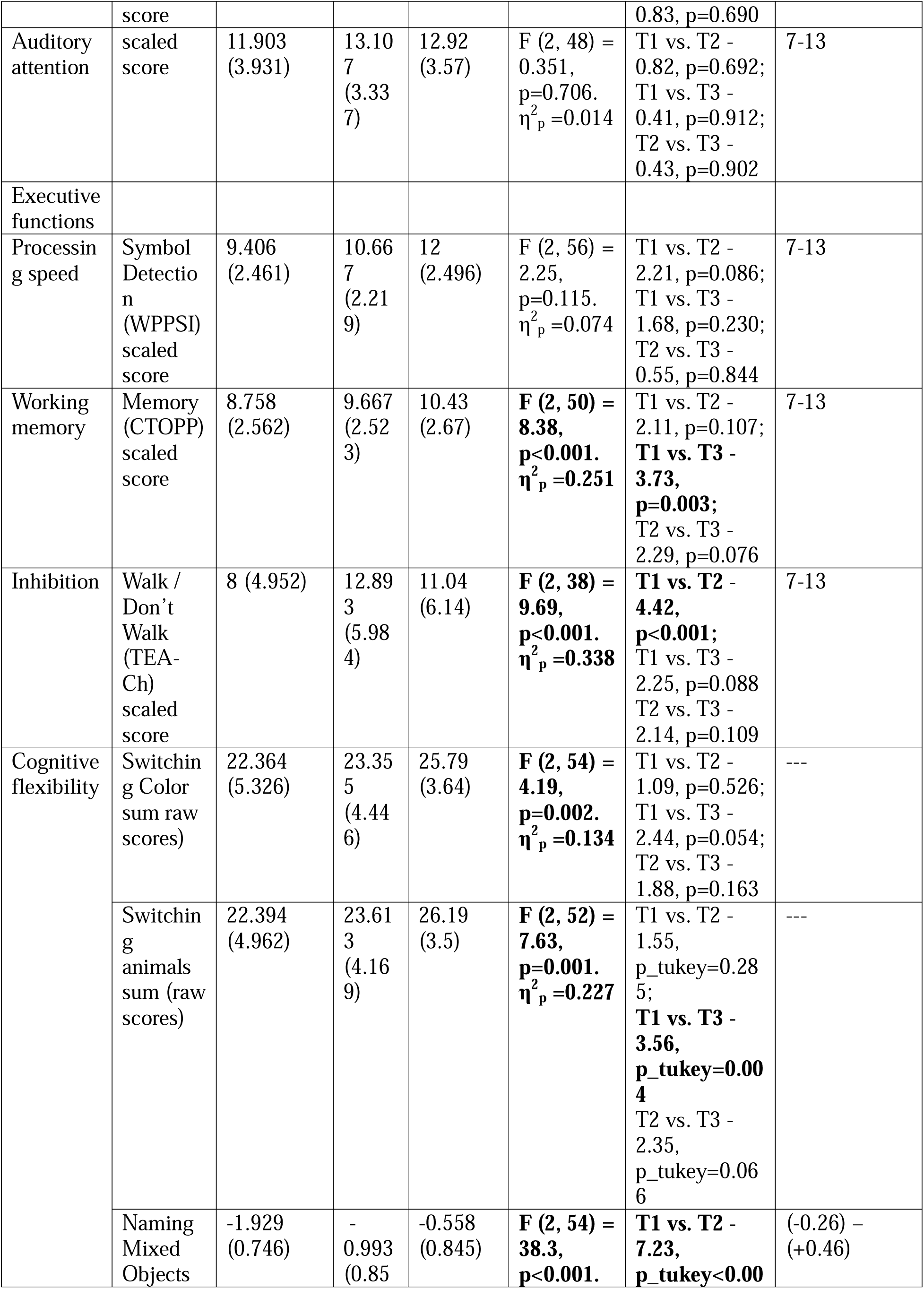
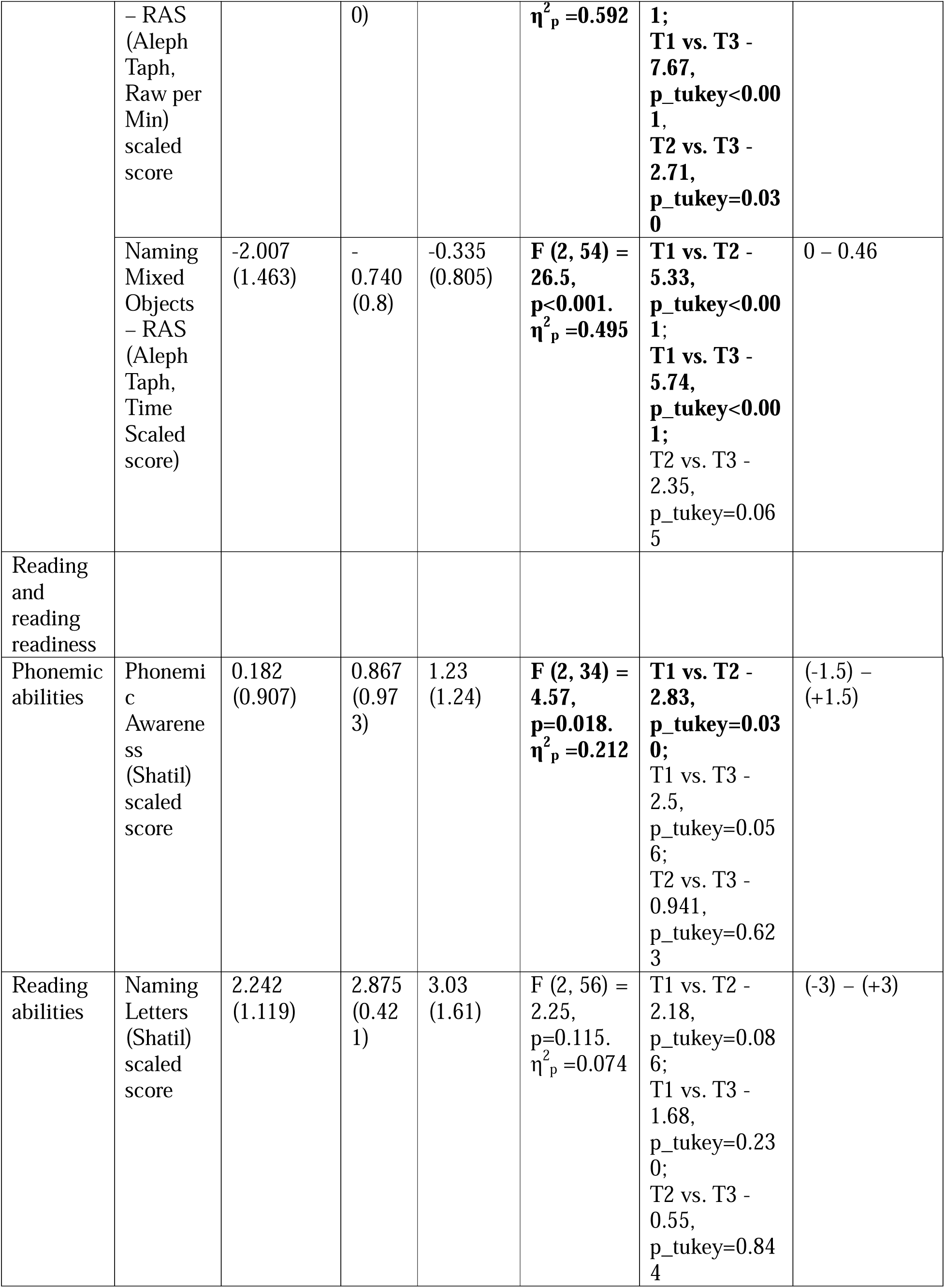
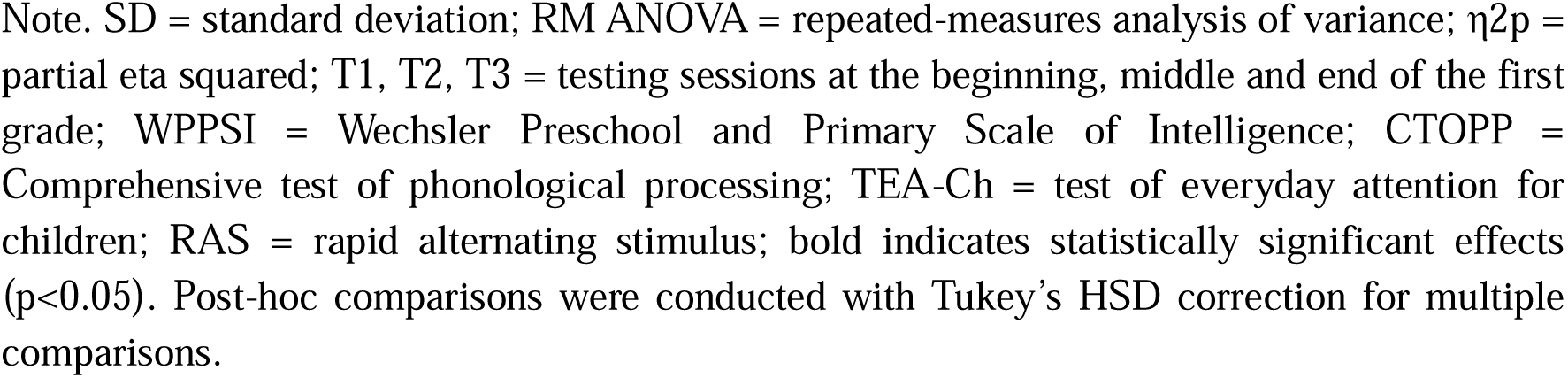
Behavioral test scores and changes from the beginning to the end of the school year.

### Accuracy and reaction times for the AVI task (outside the scanner)

Response accuracy was analyzed for 12 children who longitudinally completed the AVI task outside the scanner. Results show a significant increase in accuracy (RM ANOVA F (2,22) = 5.15, p = 0.0146) across the three sessions. Drift diffusion modeling revealed an increase in drift rate (β = 0.912, T=3.54, p=0.0001), suggesting the participants process the information faster and more accurately with further reading experience (Figure 2).

**Figure 2.**
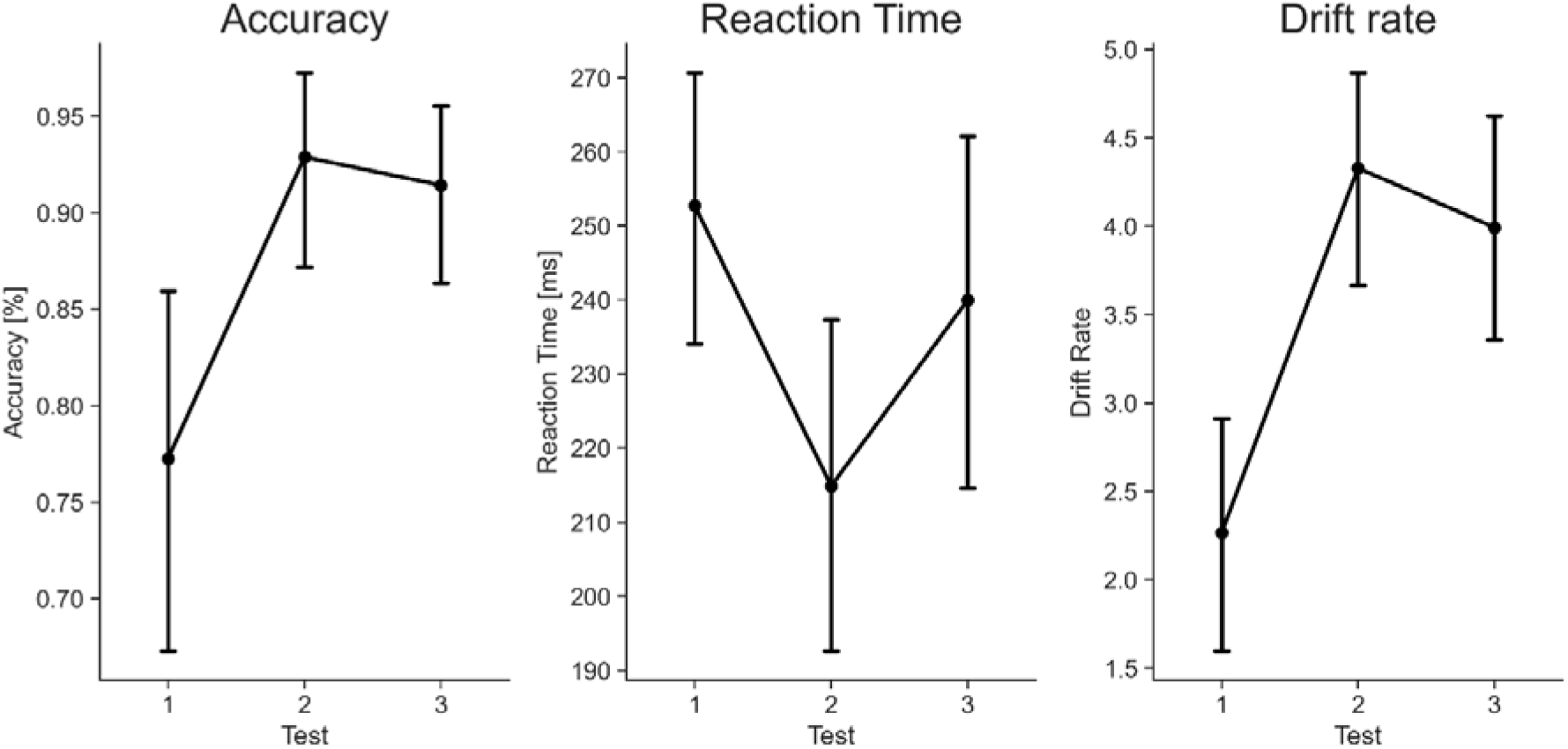
Behavioral audiovisual matching performance across the three testing sessions outside of the MRI (offline). The left and middle panels show the observed behavioral measures (accuracy and reaction time). The left panel shows the drift rate estimated using drift diffusion modeling.

### Neuroimaging results

Repeated-measures ANOVA of mean changes in hemodynamic activation (beta values) obtained from predefined regions of interest (ROIs) by estimating a general linear model revealed a significant increase in neural activation for the congruent minus incongruent contrast occurring predominantly between the middle and end of the first grade. The increase is significant only in the fusiform gyrus [F (2,42) = 5.1, p = 0.01, η^2^_p_=0.18] (both hemispheres taken together) and [F (2,42) = 4.04, p = 0.025, η^2^_p_=0.16] (right hemisphere). In the left hemisphere, the increase is significant across all defined ROIs ([frontal: F (2,42) = 3.61, p = 0.036 η^2^_p_=0.15], superior temporal gyrus: [F (2,42) = 4.73, p = 0.014, η^2^ =0.18], fusiform gyrus: [F (2,42) = 6.34, p = 0.004, η^2^ =0.23], parietotemporal: [F (2, 42) = 4.84, p = 0.013, η^2^ =0.18]). Post-hoc t-tests confirm that no significant change is observed between Test 1 and Test 2, with the significant increase appearing only from the middle to the end of the year (Figure 2).

**Figure 2:**
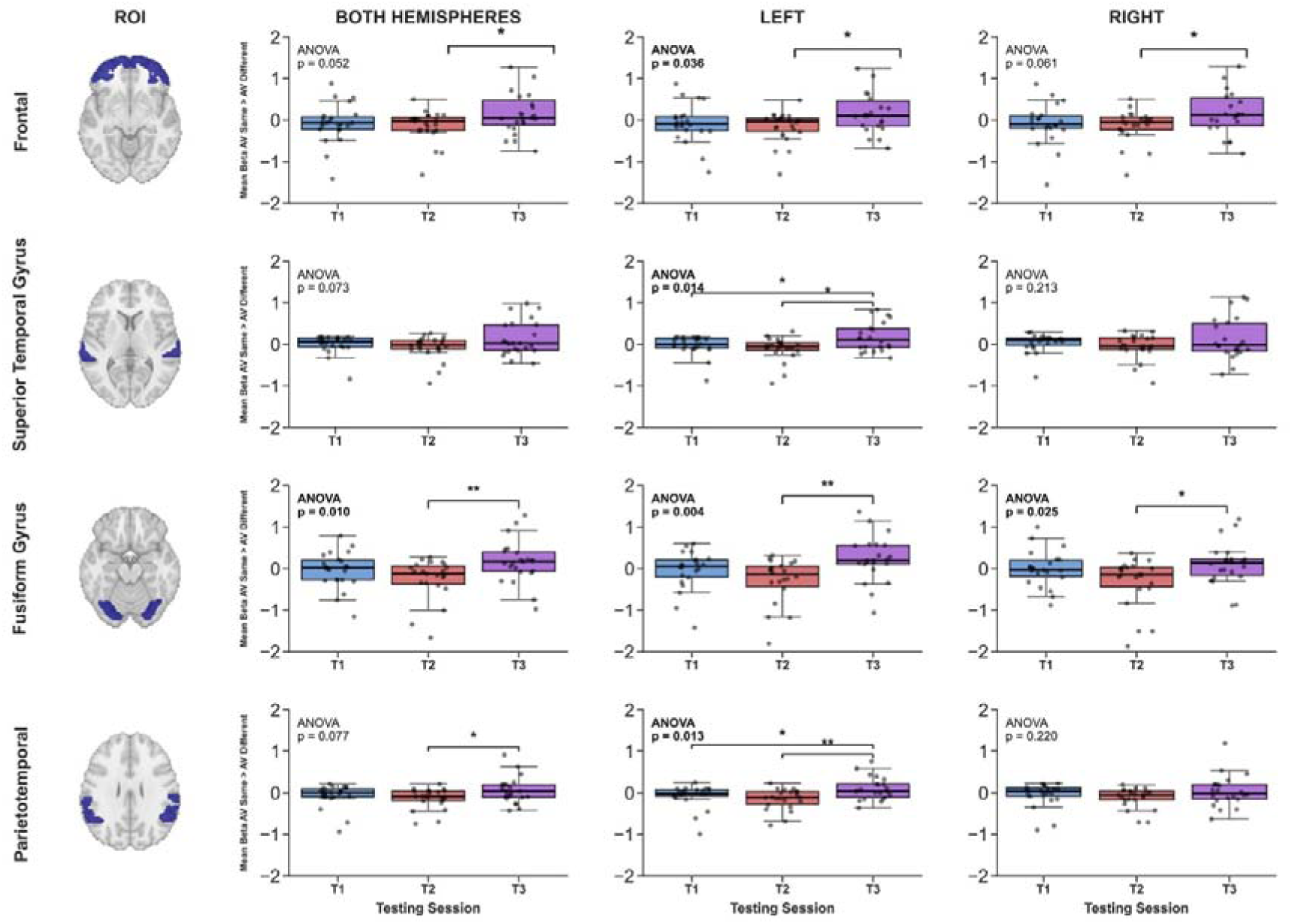
Longitudinal ROI analysis for four reading-related regions across three testing sessions in the first school year. Rows from top to bottom: frontal, superior temporal gyrus, fusiform gyrus and parietotemporal cortex. Columns from left to right: both hemispheres, left hemisphere and right hemisphere. Bars represent mean beta values at Test 1 (blue), Test 2 (orange) and Test 3 (purple). Repeated measures ANOVA results are displayed within each panel, with significant effects (p<0.05) in bold. Asterisks indicate significant post-hoc comparisons.

### Lateralization across ROIs

One-sided t-tests were used to explore the differences in activation in the left and right hemispheres for all ROIs. Results show an increase in activation in both hemispheres from the middle to the end of the year, with a significant lateralization effect in the fusiform gyrus (T (24) = 2.589, p = 0.0081) at the end of the first grade.

**Figure 3:**
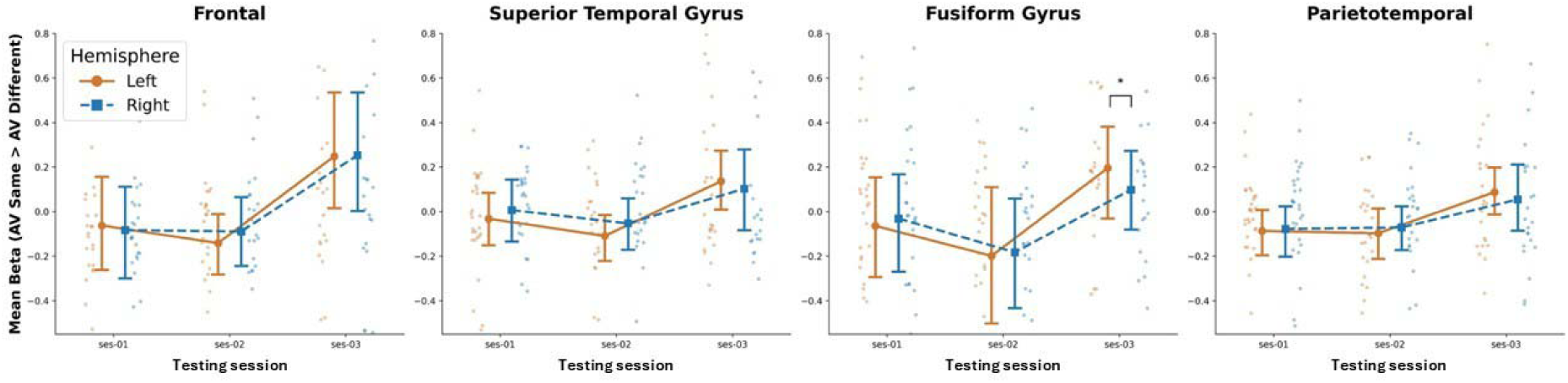
Lateralization effect across the three testing sessions in the four ROIs. The y-axis shows the mean beta value for the AV match > AV non-match contrast, extracted from the left (orange) and right (blue) hemisphere masks for each region. Points and squares represent group means and bars represent standard deviation; left- and right-hemisphere markers are offset horizontally for visibility, although data were acquired in the same sessions.

### Associations between neuroimaging and behavioral measures

To explore the relationship between behavioral measures and functional brain activation, a Linear Mixed-Effects model (LMM) was conducted to associate the standardized scores of the Naming Objects and Colors test with the extracted beta values for all ROIs. The statistical test was run using naming scores and session as a fixed effect and a by-subject random intercept to account for repeated measurements across the three sessions.

Of the four ROIs tested, only the fusiform gyrus yielded a significant positive association between naming objects and colors scores and functional activation (β=0.109, p=0.027). This indicates that during reading acquisition, higher behavioral scores are associated with increased neural activation in this region (Figure 5).

**Figure 4.**
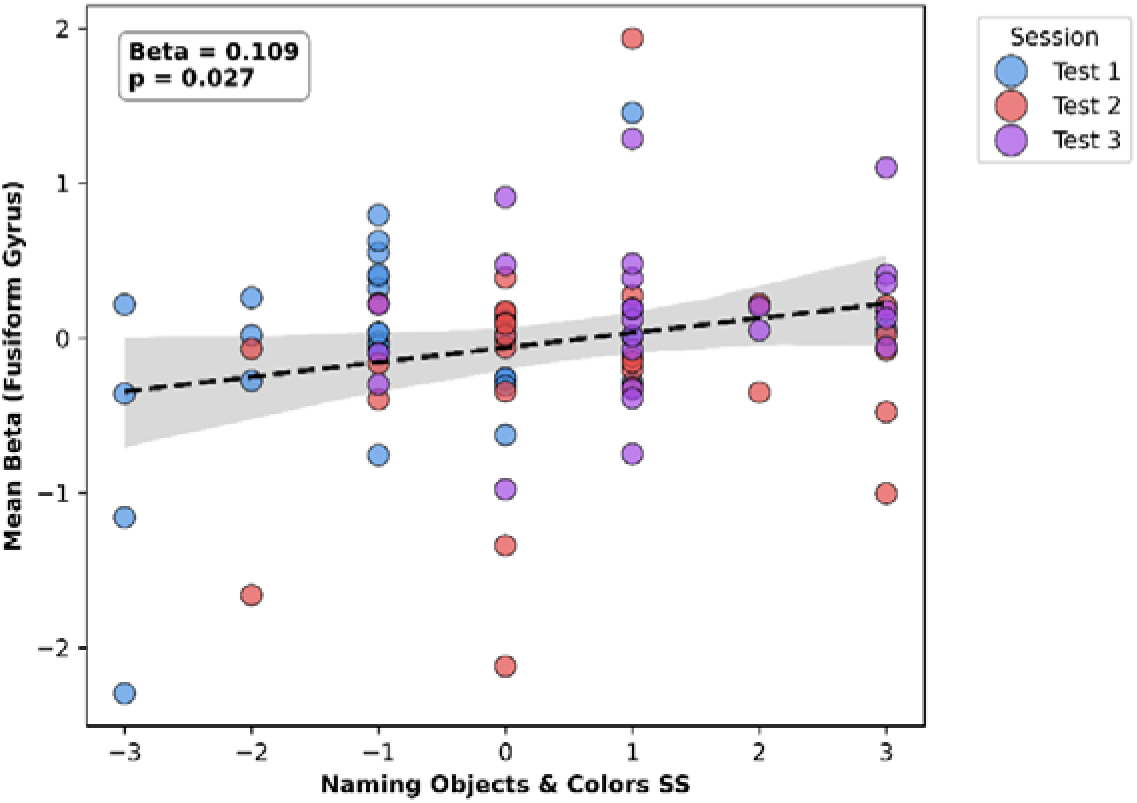
Correlation between naming objects and colors scores and fusiform gyrus activation across all sessions. X-axis: naming objects and colors scaled scores, y-axis: mean functional beta for the fusiform gyrus. Each dot represents one participant at Test 1 (blue), Test 2 (red), or Test 3 (purple).

## Discussion

The current study aimed to characterize the neural and behavioral development of AV integration processing during first-grade Hebrew reading acquisition, and to test whether this development was related to naming behavioral scores. Three key results were identified. First, the fusiform gyrus showed a significant bilateral increase in activation across the school year, while the superior temporal gyrus, frontal cortices, and parietotemporal cortices showed a significant increase restricted to the left hemisphere. Across all these regions, the neural changes were concentrated in the second half of the year, pointing to a more abrupt shift in multisensory integration rather than gradual development across first grade. Second, the bilateral increase that was stronger in the left hemisphere resulted in a significant left lateralization effect in the fusiform gyrus by the end of the year. Lastly, fusiform gyrus activation was positively correlated with naming scores across the first grade, directly linking neural development to executive function scores.

### Recruitment of audiovisual integration regions increased across the first grade

The clearest evidence of increased engagement in the AV integration regions was found for the fusiform gyrus, which showed a significant increase in activation overall and in both hemispheres. This was consistent with the fusiform gyrus’s role in housing the Visual Word Form Area (VWFA), whose primary function is fast and automatic parallel processing of letters and words (Barton et al., 2025; Kweldju, 2015), and aligned with previous findings linking increased activation in this region during letter-sound processing to growing reading fluency (Karipidis et al., 2021). Since first-grade children were still building their letter-sound mappings, increased recruitment in this region may be related to more efficient visual word-form processing developing alongside phonological processing.

We also found a significant increase in activation in the left superior temporal gyrus, frontal cortices, and parietotemporal cortices. The superior temporal gyrus is associated with the convergence of orthographic and phonological information (Gao et al., 2023), and our findings were consistent with Karipidis et al. (2021), who describe an increase in left STG activity from the pre-reading stages to fluent reading in typically developing children. Frontal regions are associated with executive control, error monitoring and attentional modulation (Li et al., 2023) and the increase in left frontal regions may reflect strengthening articulatory and phonological processing as children became more proficient at translating graphemes into phonemes (Li et al., 2023; Ren et al., 2026). The parietotemporal cortex supports reading through auditory and visual feature analysis and attention shifting (Raij et al., 2000; Ren et al., 2026). These regions did not show a bilateral increase or a significant lateralization effect, possibly because their engagement was already left-dominant or would become left-dominant in a different window of development than that included in this study.

The timing of the increased activation was also notable. Post-hoc comparisons revealed no significant changes between the beginning and the middle of the school year, with all significant increases emerging only between the middle and the end of the school year. This observation complements the findings in Karipidis et al. (2021), who describe an increase in activation in the VWFA, STG, and IFG that emerges rapidly between kindergarten and the first grade. Our data suggest the increase was concentrated in the second half of the first year rather than distributed evenly across the school year. These findings align with the alphabetic stage of Frith’s (1986) reading model, which is characterized by effortful phonological decoding. Our results suggest that the second half of the first grade may represent the window in which children have accumulated sufficient letter-to-sound knowledge and experience to begin transitioning from the alphabetic to the orthographic stage, associated with more automatic letter recognition.

### Left lateralization in the fusiform gyrus emerged at the end of the first grade

Our hypothesis regarding left lateralization was confirmed specifically for the fusiform gyrus. Although activation in this region increased significantly in both hemispheres from the beginning to the middle of the first grade, the left hemisphere outpaced the right, resulting in a significant difference between hemispheres by the end of the year. This can be understood within the developmental framework described by Sadick & Ginsburg (1978), which characterized early readers aged five to seven as ambilateral, actively engaging both hemispheres during reading-related processing. Our data reflects exactly this transitional state: both hemispheres were initially recruited, with left lateralization beginning to emerge as letter-sound knowledge accumulated across the year. Within this developmental trajectory, the right hemisphere reduction described by Seghier & Price (2011) as a mechanism of lateralization likely represents a later stage in reading development, occurring after more reading experience has consolidated orthographic representations to the point where right hemisphere engagement is no longer necessary. Our sample captured the early stages of this process, before the specialization described in older children and adult samples begins.

### Naming skills are associated with fusiform gyrus activation across the first grade

Behavioral scores improved significantly across the first year in skills related to general abilities, phonemic processing and executive functions. However, the most direct link to the neural findings was a significant positive correlation between letter naming scores and fusiform gyrus activation, confirming that neural specialization in this region tracks reading-related knowledge. The timing of improvement differed across measures. Inhibition and phonemic awareness improved predominantly between the beginning and the middle of the school year, with little additional gain towards the end of the school year, which may reflect the role of these cognitive functions as foundational prerequisites for reading. General verbal abilities followed the opposite pattern, improving predominantly between the middle and the end of the school year. Working memory and cognitive flexibility followed a third pattern of steady increase across the three timepoints, reflecting their nature as domain-general skills that develop gradually. Overall, these findings suggested that different cognitive and pre-literacy skills did not develop uniformly.

The performance in the AV integration task itself showed a clear improvement across the year, with both accuracy and drift rate increasing significantly. The rise in drift rate indicated that children were not only more accurate but also accumulating information faster and more efficiently when matching auditory and visual stimuli, consistent with faster and more automatic processing as the year progressed. The timing suggested that the improvement occurred predominantly between the beginning and middle of the first grade, with little behavioral gain by the end of the year. This was different from our neuroimaging results, where the significant increase in activation occurred almost exclusively between the middle and end of the year. This dissociation suggests that behavioral task performance and neural specialization follow distinct developmental timelines during early reading acquisition.

Basic letter-sound associations, sufficient for correct behavioral performance on a matching task, may develop earlier than the neural sensitivity to the distinction between congruent and incongruent letter-sound pairs that the fMRI contrast captures. Children may have achieved good task performance before the brain had fully developed the specificity required to discriminate letter-sound congruency at a neural level, with this more refined tuning developing in the second half of the year.

Our results showed a significant positive correlation between naming objects and colors scores and hemodynamic activation in the fusiform gyrus, the region showing the strongest developmental increase, confirming our third hypothesis and highlighting a neurocognitive link during early reading acquisition. The naming objects and colors rapid alternating stimulus (RAS) challenges the child’s ability to connect visual stimuli to verbal labels, demanding high levels of cognitive flexibility, rapid memory retrieval, and efficient semantic processing (Albuquerque & Simões, 2010). At a neural level, the fusiform gyrus develops rapidly as children transition to fluent reading (Horowitz Kraus, 2023; Joseph et al., 2001; McCandliss et al., 2003). During the first grade, children are transitioning from effortful phonological decoding to automatic letter-sound recognition. Therefore, this positive correlation suggests that children with stronger skills in rapid visual-verbal integration are more efficient in recruiting the fusiform gyrus during naming tasks. This relationship further carries clinical relevance because both reduced fusiform gyrus activation and low naming scores are established hallmarks for developmental dyslexia (Albuquerque & Simões, 2010; Horowitz Kraus, 2023). Their coupled developmental trajectory suggests a shared vulnerability. The intersection between poor RAS performance and delayed fusiform gyrus recruitment during the first grade could potentially serve as a biomarker for reading disabilities, allowing for early identification and intervention in vulnerable children.

### Limitations

This study used a priori ROIs based on known key regions that contribute to AV integration according to the literature. This approach allowed us to test specific hypotheses about the development of these regions. However, it also means that our analysis was restricted to these predefined regions and we might be missing other developmental changes occurring outside these ROIs. The ROI analyses involved multiple comparisons across four regions and three hemisphere masks (bilateral, left and right) without correction, therefore the reported results should be interpreted with caution. Additionally, no motion-based exclusion for subjects or sessions was applied given the expected elevated head motion in this pediatric sample. The effect of head motion was instead addressed by regressing out motion indices when estimating the first-level statistical model. Furthermore, drift-diffusion modeling (DDM) analysis was conducted on a subset of participants (n=12) as a systematic collection of behavioral performance data outside the scanner was implemented partway through the study. Accordingly, direct correspondence between the DDM results and the neuroimaging data should be interpreted with caution. For future investigations, we propose evaluating the role of the environment in these developmental trajectories. External factors, such as socioeconomic status, parental stress, or war-related stress, likely have an effect on the developing brain. Understanding how these environmental inputs interact with the development of executive functions and AV congruency processes is crucial to get a complete picture of reading acquisition, guiding to more tailored interventions for children at risk of reading difficulties.

### Conclusions

In conclusion, these findings suggest that the fusiform gyrus emerged as the central region of audiovisual congruency processing, showing bilateral engagement, progressive left lateralization, and a direct relationship with RAS naming skills. Additionally, the second half of the first grade turned out to be a critical window for the neural specialization underlying letter-sound mapping, distinct from the earlier window in which behavioral competence on the audiovisual task was established. Together, these findings contribute to a deeper understanding of how audiovisual integration develops during the first year of reading acquisition.

## Acknowledgments

This study was supported by the Deutsche Forschungsgemeinschaft (DFG, German Research Foundation, Grant Number 499339749).

## References

Abraham, A., Pedregosa, F., Eickenberg, M., Gervais, P., Mueller, A., Kossaifi, J., Gramfort, A., Thirion, B., & Varoquaux, G. (2014). Machine learning for neuroimaging with scikit-learn [Methods]. Frontiers in Neuroinformatics, Volume 8 - 2014. 10.3389/fninf.2014.00014

Albuquerque, C. P., & Simões, M. R. (2010). Rapid naming tests: Developmental course and relations with neuropsychological measures. The Spanish journal of psychology, 13(1), 88–100.

Barton, J. J., Albonico, A., & Starrfelt, R. (2025). The lateralization of reading. Handbook of Clinical Neurology, 208, 301–325.

Esteban, O., Markiewicz, C. J., Blair, R. W., Moodie, C. A., Isik, A. I., Erramuzpe, A., Kent, J. D., Goncalves, M., DuPre, E., & Snyder, M. (2019). fMRIPrep: a robust preprocessing pipeline for functional MRI. Nature methods, 16(1), 111–116.

Frith, U. (1986). A developmental framework for developmental dyslexia. Annals of dyslexia, 36(1), 67–81.

Gao, C., Green, J. J., Yang, X., Oh, S., Kim, J., & Shinkareva, S. V. (2023). Audiovisual integration in the human brain: a coordinate-based meta-analysis. Cerebral Cortex, 33(9), 5574–5584.

Holloway, I. D., van Atteveldt, N., Blomert, L., & Ansari, D. (2015). Orthographic dependency in the neural correlates of reading: evidence from audiovisual integration in English readers. Cerebral Cortex, 25(6), 1544–1553.

Horowitz-Kraus, T., Cancer, A., Antonietti, A., Rosch, K., & Farah, R. (2025). Audiovisual integration and cognitive control supporting reading fluency. Advances in child development and behavior, 68, 83–104.

Horowitz-Kraus, T. (2023). The role of executive functions in fluent reading: Lessons from reading acquisition and remediation. Mind, Brain, and Education, 17(4), 373–382.

IBM. IBM SPSS Statistics. In https://www.ibm.com/products/spss-statistics

Jamovi. (2024). The Jamovi Project. In https://www.jamovi.org

Joseph, J., Noble, K., & Eden, G. (2001). The neurobiological basis of reading. Journal of Learning Disabilities, 34(6), 566–579.

Karipidis, I. I., Pleisch, G., Di Pietro, S. V., Fraga-González, G., & Brem, S. (2021). Developmental trajectories of letter and speech sound integration during reading acquisition. Frontiers in psychology, 12, 750491.

Kweldju, S. (2015). Neurobiology Research Findings: How the Brain Works during Reading. Pasaa: Journal of language teaching and learning in Thailand, 50, 125–142.

Lenth, R. (2023). emmeans: Estimated Marginal Means, aka Least-Squares Means_. R package version 1.8. 5.

Li, J., Yang, Y., Viñas-Guasch, N., Yang, Y., & Bi, H. Y. (2023). Differences in brain functional networks for audiovisual integration during reading between children and adults. Annals of the New York Academy of Sciences, 1520(1), 127–139.

Manly, T., Anderson, V., Nimmo-Smith, I., Turner, A., Watson, P., & Robertson, I. H. (2001). The differential assessment of children’s attention: The Test of Everyday Attention for Children (TEA-Ch), normative sample and ADHD performance. The Journal of Child Psychology and Psychiatry and Allied Disciplines, 42(8), 1065–1081.

McCandliss, B. D., Cohen, L., & Dehaene, S. (2003). The visual word form area: expertise for reading in the fusiform gyrus. Trends in cognitive sciences, 7(7), 293–299.

Myers, C. E., Interian, A., & Moustafa, A. A. (2022). A practical introduction to using the drift diffusion model of decision-making in cognitive psychology, neuroscience, and health sciences. Frontiers in psychology, 13, 1039172.

Pleisch, G., Karipidis, I. I., Brauchli, C., Röthlisberger, M., Hofstetter, C., Stämpfli, P., Walitza, S., & Brem, S. (2019). Emerging neural specialization of the ventral occipitotemporal cortex to characters through phonological association learning in preschool children. NeuroImage, 189, 813–831.

Power, J. D., Barnes, K. A., Snyder, A. Z., Schlaggar, B. L., & Petersen, S. E. (2012). Spurious but systematic correlations in functional connectivity MRI networks arise from subject motion. NeuroImage, 59(3), 2142–2154.

Pstnet, I. (2020). E-Prime 3.0. in Psychology Software Tools. In https://pstnet.com/products/e-prime/

Raij, T., Uutela, K., & Hari, R. (2000). Audiovisual integration of letters in the human brain. Neuron, 28(2), 617–625.

Ren, J., Wang, H., Landi, N., Joanisse, M. F., Gracco, V., Kleinman, D., Mahaffy, K., Bajracharya, A., Hancock, R., & Pugh, K. (2026). Audiovisual integration in reading among school-aged children: Evidence from combined fMRI and EEG. Developmental Cognitive Neuroscience, 101729.

Sadick, T. L., & Ginsburg, B. E. (1978). The development of the lateral functions and reading ability. Cortex, 14(1), 3–11.

Seghier, M. L., & Price, C. J. (2011). Explaining left lateralization for words in the ventral occipitotemporal cortex. Journal of Neuroscience, 31(41), 14745–14753.

Shany, M., Lachman, D., Shalem, Z., Bahat, A., & Zeiger, T. (2006). Alef-Taf, Evaluation system for the diagnosis of disability in the reading and writing process according to national norms. Holon: Yesod Publishing.(Hebrew).

Shatil, E. (2000). Shatil test for the assesment of early childhood literacy. Israel: Ach Publishers.

Shatil, E., & Share, D. L. (2003). Cognitive antecedents of early reading ability: A test of the modularity hypothesis. Journal of experimental child psychology, 86(1), 1–31.

Shaywitz, S. E., & Shaywitz, B. A. (2007). The neurobiology of reading and dyslexia. The ASHA Leader, 12(12), 20–21.

Singmann, H., Bolker, B., Westfall, J., Aust, F., & Ben-Shachar, M. S. (2012). afex: Analysis of factorial experiments. (No Title).

Team, R. C. (2024). A Language and environment for statistical computing. In https://cran.r-project.org/

Wagner, R. K., Torgesen, J. K., Rashotte, C. A., & Pearson, N. A. (1999). Comprehensive test of phonological processing: CTOPP. Pro-ed Austin, TX.

Wechsler, D. (2012). Wechsler preschool and primary scale of intelligence—fourth edition. The Psychological Corporation San Antonio, TX.

Ziv, Y. (2017). Animals and colors inhibition/switching test. *Unpublished Test* (Haifa: University of Haifa, 2017).

